# SynFit: Synergistic Contrastive Learning for Multi-Objective Protein Fitness Prediction and Optimization

**DOI:** 10.64898/2026.05.21.726972

**Authors:** Tony Tu, Wei Huang, Ziang Li, Kerr Ding, Yang Yang, Yunan Luo

## Abstract

Proteins function through a complex interplay of structural and biochemical properties, and mutations can reshape these properties to generate fitness landscapes spanning multiple functional objectives. A central challenge in protein engineering is the need to simultaneously optimize multiple properties. In biocatalysis, for example, practical enzyme development routinely requires the concurrent optimization of catalytic activity, selectivity, stability, and substrate generality. However, despite recent advances in computational protein design and fitness prediction, most existing approaches treat these properties independently and do not explicitly capture the dependencies and trade-offs that govern real-world protein performance. We present SynFit, a multi-objective learning framework that integrates pretrained protein language models with experimental fitness measurements for protein fitness prediction and engineering. SynFit learns both shared and property-specific protein sequence representations through a synergistic contrastive learning strategy, enabling the identification of variants that simultaneously optimize multiple functional properties. Across a large-scale multi-fitness deep mutational scanning benchmark, SynFit consistently outperforms state-of-the-art supervised models trained on individual objectives and more accurately identifies variants that balance competing functional constraints. We further applied SynFit to multi-objective enzyme design for a new-to-nature biocatalytic enantioselective borylation reaction, providing a diverse array of novel cytochrome *c* sextuple variants in a single round of design with simultaneously improved catalytic activity and enantioselectivity that rival the best variants obtained through directed evolution. Together, these results establish SynFit as a general framework for multidimensional protein fitness prediction and highlight its potential to enable efficient multi-objective optimization in protein engineering, particularly in biocatalysis.

## 1 Introduction

Predicting the functional consequences of amino acid substitutions remains a fundamental challenge in protein science and engineering. Traditional computational approaches, including physics-based energy models and supervised machine learning (ML) methods, have provided important foundations for mutational effect prediction, but they often rely on engineered features, assay-specific supervision, and sufficiently large labeled datasets ^1–7^. More recently, pretrained protein language models (pLMs) have substantially expanded the scope of sequence-to-fitness prediction by learning evolutionary and structural constraints from large protein sequence databases, enabling zero-shot predictions across diverse proteins and functions ^8–15^. However, because these models primarily capture patterns of natural sequence variation rather than task-specific experimental readouts, their zero-shot predictions generally remain less accurate than models adapted to function-specific measurements ^16–21^. In this context, integrating pretrained representations with limited experimental data through fine-tuning represents a highly appealing approach to bridging general sequence knowledge with function-specific prediction ^17;21–26^.

Despite recent progress, most existing approaches remain focused on single-property prediction, whereas protein function is inherently multi-objective. In practice, successful protein engineering requires the simultaneous optimization of multiple functional properties. For example, in the field of biocatalysis and enzyme engineering, the development of useful biocatalysts often requires concurrent optimization of catalytic activity, selectivity, stability, and substrate generality. More broadly, mutations frequently affect stability, catalytic efficiency, binding affinity, expression, and selectivity in coupled and context-dependent ways ^27–31^. These interdependencies give rise to complex fitness landscapes that cannot be readily decomposed into independent objectives. This problem is further exacerbated by the small and heterogeneous nature of experimental datasets, particularly in the low-data regime where models must disentangle shared sequence features from property-specific determinants. Collectively, these challenges highlight the need for methods that integrate pretrained sequence knowledge with limited experimental data while explicitly capturing dependencies and trade-offs across multiple functional properties.

Here, we introduce SynFit (<u>syn</u>ergistic contrastive learning for <u>fi</u>tness prediction), a deep learning framework that integrates pretrained pLMs with experimental fitness measurements for multi-objective protein fitness prediction and engineering. SynFit combines a shared module that learns sequence features informative across diverse functions and a set of property-specific predictors that exploit the learned shared features for accurate multi-objective fitness prediction. To address the challenges of data scarcity, SynFit introduces a synergistic contrastive learning algorithm that effectively recalibrates pretrained pLM representations toward assay-specific fitness data while avoiding overfitting. Based on these predictions, SynFit prioritizes protein variants that simultaneously optimize multiple functional properties.

To evaluate SynFit, we curated a multi-fitness benchmark derived from ProteinGym ^16^, comprising 46 deep mutational scanning (DMS) assays across 20 proteins and spanning diverse functions. Across this benchmark, SynFit consistently outperforms single-objective models trained on individual assays, identifies variants that balance multiple functional constraints, and reveals residues with shared functional importance across assays. We further validated SynFit experimentally in a new-to-nature biocatalytic borylation system catalyzed by a heme-dependent cytochrome *c* protein, demonstrating that SynFit-guided designs could furnish a diverse collection of novel combinatorial biocatalyst variants with catalytic activity and enantioselectivity that rival or surpass those obtained through laboratory directed evolution. Sequence analysis of top-performing variants revealed convergent residue preferences across the engineered active-site positions, consistent with a focused, multi-objective optimization of the combinatorial sequence space. Together, these results establish SynFit as a general framework for multi-objective protein fitness prediction and engineering and highlight its potential to advance ML–guided protein engineering, particularly for use in biocatalysis.

## 2 Results

### 2.1 Synergistic contrastive learning for multi-objective fitness prediction

Protein engineering often requires optimizing multiple functional properties simultaneously, such as activity, stability, or selectivity. These properties are frequently coupled: a mutation that improves one aspect of function may weaken another, whereas some mutations influence multiple properties through shared structural or biochemical mechanisms. In typical deep mutational scanning (DMS) experiments, however, each assay captures only one facet of protein properties, and different assays provide only partial and heterogeneous measurements of the same sequence landscape. To address this challenge, we developed SynFit, a framework that jointly learns from multiple functional assays to model the multidimensional fitness landscape of a protein (Fig. 1a).

**Figure 1:**
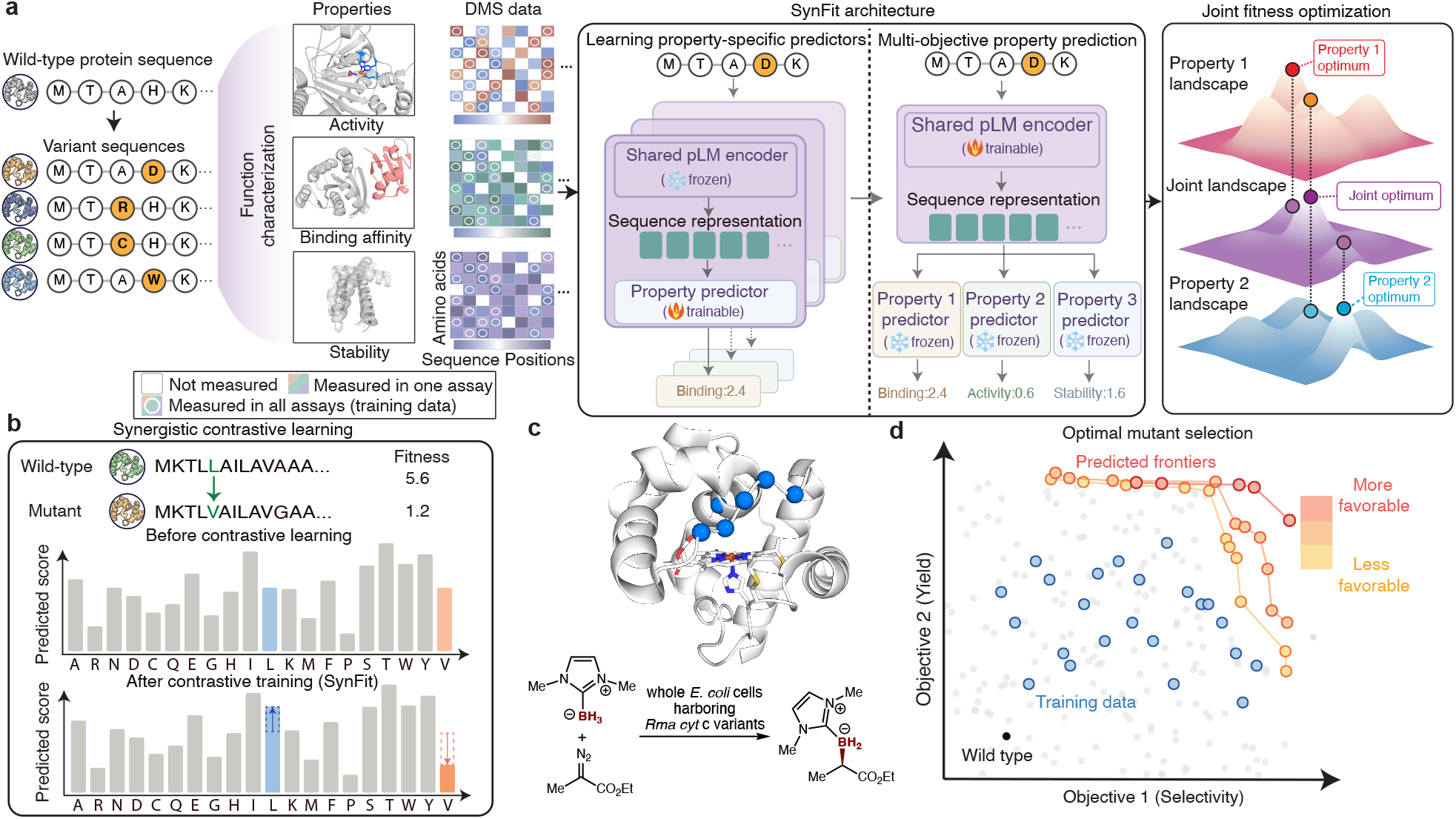
Overview of our multi-fitness prediction framework. **(a)** Protein variants derived from the wildtype sequence are experimentally characterized for a subset of properties to generate partial deep mutational scanning (DMS) data used for model training.SynFit jointly fine-tunes a pretrained protein language model across multiple assays using a shared encoder and property-specific predictors. **(b)** Each objective combines a contrastive BT loss with KL regularization against the ESM2 prior, and calibrated predictions are integrated via non-dominated sorting to yield multi-objective Pareto surfaces. **(c)** To validate SynFit, we applied it to the combinatorial optimization of *Rma* cyt *c* variants for asymmetric carbene B–H insertion, a new-to-nature biocatalytic reaction that converts NHC·BH_3_ and *α*-diazocarbonyl compounds to chiral organoboron products in whole *E. coli* cells. **(d)** SynFit-predicted multi-objective fitness landscape for 909 experimentally characterized *Rma* cyt *c* variants, with Pareto-optimal candidates selected for experimental validation.

SynFit starts from a pretrained pLM, which encodes broad evolutionary information from natural protein sequences. Building on this backbone architecture, SynFit learns a shared representation that captures sequence features relevant across functional readouts, together with property-specific predictors that model individual objectives (Fig. 1b). In this way, the framework captures both common constraints that shape multiple biochemical properties and the distinct signals associated with each assay, rather than treating each measurement independently. We experimentally validated SynFit on the combinatorial optimization of *Rma* cyt *c* variants for asymmetric carbene B–H insertion (Fig. 1c), a new-to-nature biocatalytic reaction optimized jointly for yield and enantioselectivity. The resulting predictions can then be used to prioritize variants that achieve the best overall balance across multiple objectives, which is essential for practical protein engineering applications (Fig. 1d).

This framework is designed to make multi-objective learning effective even when experimental data are limited. Rather than extensively retraining the pretrained model to fit absolute assay values, SynFit adapts it in a data-efficient manner by learning relative differences among variants within each assay. This strategy builds on our previous low-*N* fitness prediction model ^23^ while advancing it to the multi-objective setting through joint learning across multiple functional assays. This strategy preserves the broad biological information encoded during pretraining while recalibrating the model toward experimentally measured function, thereby reducing overfitting to small and noisy datasets. Together, these design features enable SynFit to learn robust and biologically meaningful multidimensional fitness landscapes from sparse multi-assay measurements.

### 2.2 SynFit enables accurate and generalizable protein fitness prediction

To evaluate whether the multi-objective learning of SynFit improves protein fitness prediction, we curated a comprehensive benchmark of proteins with multiple DMS assays from ProteinGym ^16^. Specifically, we combined individual single-property functional assays from various studies characterizing the same protein and retained overlapping single-mutation variants with quantitative measurements across all available assays, constructing a “multi-fitness” dataset for rigorous evaluation of multi-objective prediction. The resulting benchmark includes 46 assays across 20 proteins spanning diverse organisms, with each protein annotated by 2-4 functional properties (Table S1). This dataset enables a systematic assessment of multi-objective fitness prediction performance.

We first tested whether synergistic learning across assays improves predictive accuracy compared to models trained on individual properties. SynFit was trained jointly on all available assays for each protein and evaluated on individual objectives, alongside established single-objective baselines. Across proteins, SynFit achieves the best performance on the majority of cases, demonstrating robustness across diverse sequence families (Fig. 2a). When aggregated across all assays, SynFit attains the highest mean and median Spearman correlations under five-fold cross-validation, outperforming strong baselines such as ConFit ^23^ and ProteinNPT ^32^ (Fig. 2b). These results indicate that multi-objective modeling across related functional measurements improves generalization and enables more accurate prediction of mutational effects across heterogeneous biochemical contexts.

**Figure 2:**
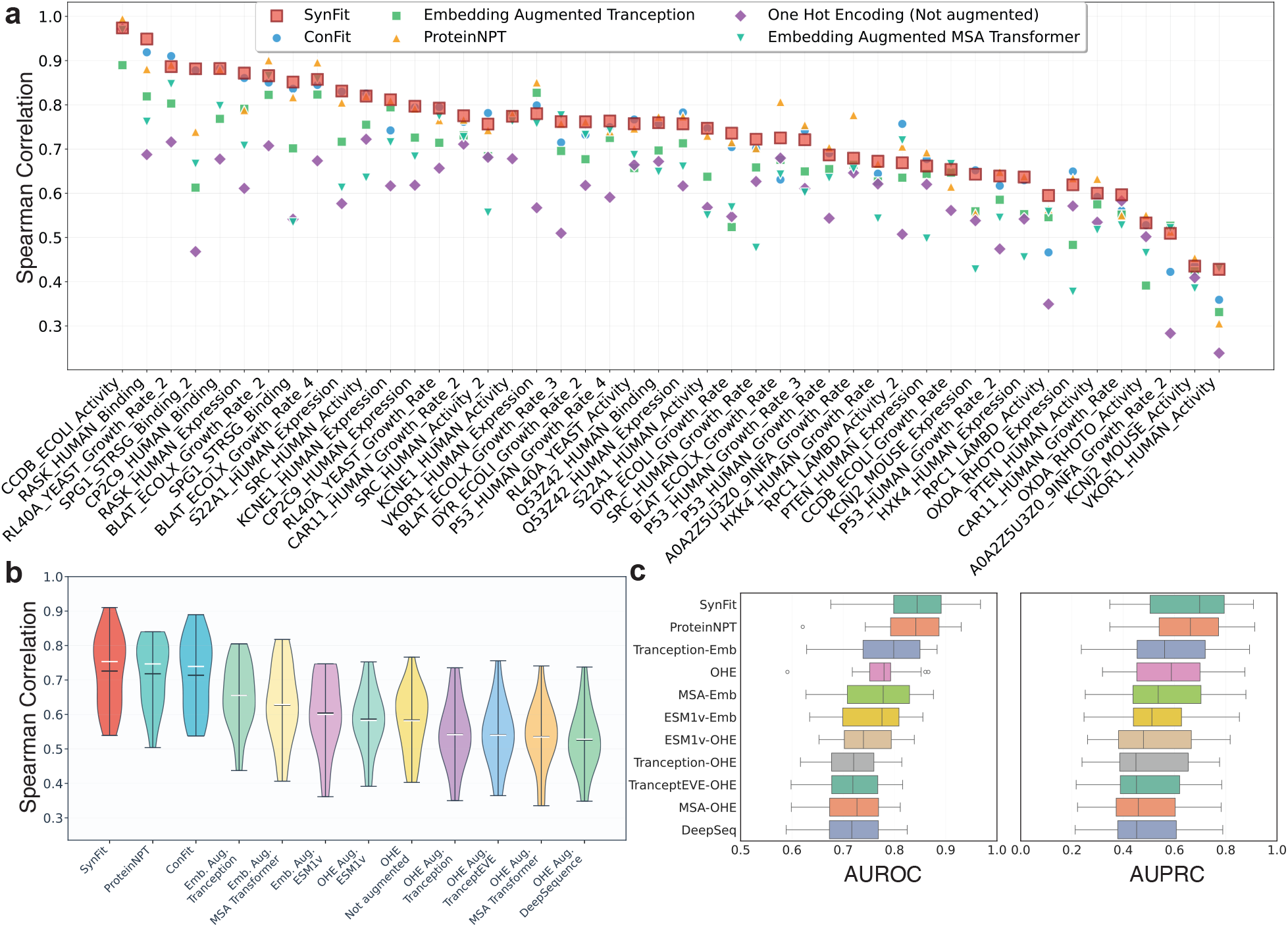
Comprehensive evaluation of SynFit function prediction performance across proteins and fitness assays. *Abbreviations:* OHE: one-hot encoding; emb.: embedding. **(a)** Five-fold cross-validation performance son held-out test data for top-performing methods, where SynFit demonstrates both high mean correlations and low variance, highlighting its robustness and generalizability across fitness dimensions. Each model was trained and evaluated over five random splits, and the reported values represent the average Spearman correlation between predicted and experimentally measured fitness across held-out test sets. **(b)** AUROC (left) and AUPRC (right) distributions on 13 proteins (28 DMS) with available wild-type anchors. SynFit achieves the highest overall performance, showing strong ability to identify beneficial mutants relative to wild type. **(c)** Distribution of protein-level Spearman correlations across the multi-fitness benchmark. SynFit achieves the highest average correlation across diverse proteins compared to all baselines.

Beyond correlation-based evaluation, a key objective in protein engineering is to identify variants that improve upon the wild type. To assess this capability, we evaluated models on 13 proteins labeled with explicit wild-type fitness values, which allows mutations to be binarized as beneficial or deleterious. SynFit consistently achieves the highest AUROC and AUPRC across proteins (Fig. 2c), indicating superior ability to prioritize beneficial variants from large mutational libraries.

Together, these results demonstrate that SynFit not only improves fitness prediction accuracy but also enhances the identification of functionally improved variants. By integrating complementary signals across multiple functional assays, the model more effectively captures sequence–function relationships that generalize across diverse sequence contexts.

### 2.3 SynFit identifies mutations that co-improve multiple properties

Models trained on individual properties often prioritize variants that perform well on one objective but poorly on others, limiting their ability to identify solutions that satisfy multiple functional constraints. To evaluate whether SynFit captures true cross-objective relationships, we assessed its ability to recover variants that simultaneously optimize multiple properties.

We ranked variants based on their joint performance across objectives using non-dominated sorting, which organizes variants into Pareto fronts (Supplementary Note A.1). In this sorting process, variants are grouped such that those on the first Pareto front are not outperformed by any other variant when all properties are considered together. We then quantified performance using a hit-rate metric, defined as the overlap between the top 100 variants predicted by the model and the top 100 variants identified from experimental measurements. SynFit achieves the highest average hit rate across all proteins (Fig. 3a), indicating that joint learning enables more accurate identification of variants that co-optimize multiple properties than approaches based on independently trained models.

**Figure 3:**
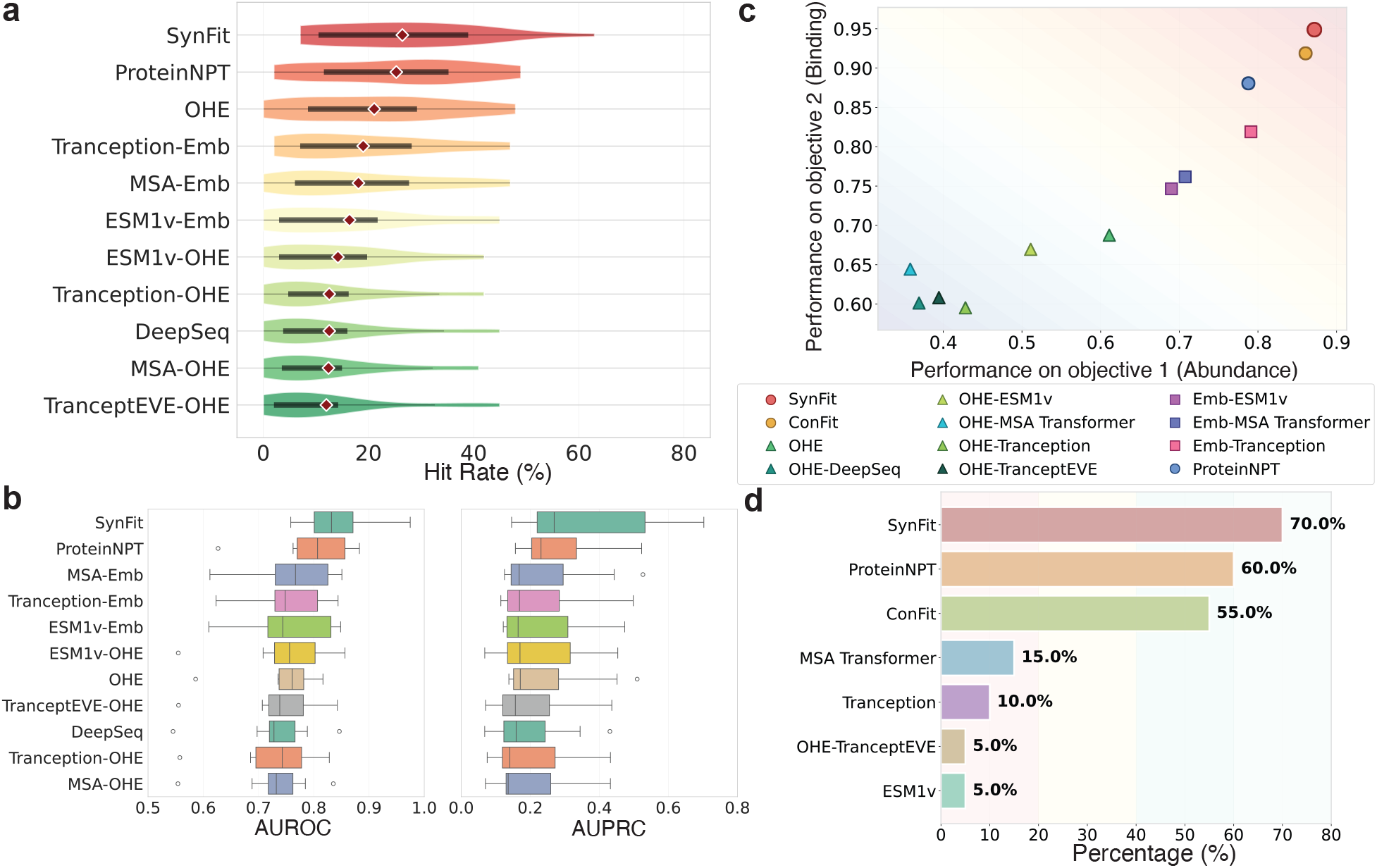
Multi-objective benchmarking of SynFit reveals synergistic optimization across fitness assays. *Abbreviations:* OHE: one-hot encoding. **(a)** Hit rate comparison of top 100 predicted versus ground-truth Pareto-optimal mutants, where each overlapping variant is counted as a “hit”, with SynFit achieving the highest overlap. **(b)** SynFit excels at identifying variants that improve upon wildtype across multiple fitness objectives, consistently enriching for jointly beneficial mutants. **(c)** Pareto analysis of held-out functional assays for *RASK_HUMAN*, based on averaged Spearman correlations across five test folds, where SynFit occupies the optimal front and outperforms all baselines. **(d)** Comparative Pareto analysis across proteins, using averaged Spearman correlations from five test folds; SynFit most frequently occupies the most optimal front.

We next evaluated the ability of SynFit to identify jointly beneficial variants, defined as mutations that outperform the wild type across all measured objectives (Fig. 3b). This represents a stringent benchmark, as variants must improve all properties simultaneously. Across eight proteins spanning 16 DMS assays with available wild-type annotations (Table S1), SynFit consistently achieves higher AUROC and AUPRC than single-objective baselines, demonstrating improved ability to prioritize variants with multi-property gains.

To further illustrate SynFit’s multi-objective capability, we analyzed the *RASK_HUMAN* dataset (Fig. 3c), which includes abundance and binding assays. Models were evaluated based on their ability to achieve optimal trade-offs among these properties. SynFit is the only model that lies on the first Pareto front (Supplementary Note A.1), indicating that it successfully captures the joint effects of mutations on both abundance and binding. Extending this analysis across all proteins, SynFit appears on the first Pareto front for 70% of proteins, outperforming ProteinNPT (60%) and ConFit (55%) (Fig. 3d). These results further support that SynFit achieves both strong per-assay accuracy while capturing shared functional representations that provide Pareto-optimal predictions across diverse protein fitness landscapes.

### 2.4 In silico case study: SynFit identifies shared functional residues in KRAS across multiple binding partners

Proteins that interact with multiple binding partners often rely on a subset of residues that mediate binding across partially overlapping interfaces. Mutations at these sites can simultaneously alter multiple interactions, thereby reshaping protein interaction networks and functional activity. Identifying such shared interaction determinants is important for understanding mutational effects and for designing proteins with controlled interaction profiles. However, models trained on individual assays typically capture only binding partner-specific signals and cannot distinguish residues that broadly regulate multiple interactions. We thus tested whether SynFit’s multi-assay learning framework can more effectively identify residues that govern binding across multiple partners. This setting naturally calls for a multi-objective approach that integrates information across assays to uncover shared biophysical constraints beyond what can be learned from any single assay.

Mutational effects often arise through distinct mechanisms. Some mutations primarily affect folding or stability, whereas others directly perturb active or interaction sites. Disentangling these effects is challenging when each assay provides only a single view of protein function ^34^. Our previous studies ^26;34^ addressed this issue by quantifying mutation effects on function beyond what can be explained by stability alone, thereby identifying functional residues. Here, SynFit extends this principle to the multi-assay setting, where multiple functional measurements provide complementary information about shared and context-specific determinants. By jointly learning across assays, SynFit can identify residues whose effects are consistently functional across different interaction contexts.

To assess whether SynFit can learn shared functional determinants across related interaction contexts, we applied it to KRAS, a central signaling protein that interacts with multiple downstream effector proteins (Fig. 4a). Recent deep mutational scanning (DMS) experiments profiled KRAS variants across six binding partners, with each assay quantitatively measuring how mutations affect binding affinity to a specific effector ^33^. Because these assays probe overlapping interaction interfaces, a unified framework should better capture shared biophysical constraints than partner-specific models. We therefore trained SynFit jointly across all six DMS datasets and compared it with individual ConFit models trained separately on each binding partner (Fig. 4b). Final residue-level function scores were averaged across independent runs and evaluated by their ability to recover known binding and allosteric sites using AUPR (Methods). Interface sites were defined by a 5 Å cutoff in PDB structures, and residues shared across all six partners were treated as common sites and used as the ground-truth positive samples.

**Figure 4:**
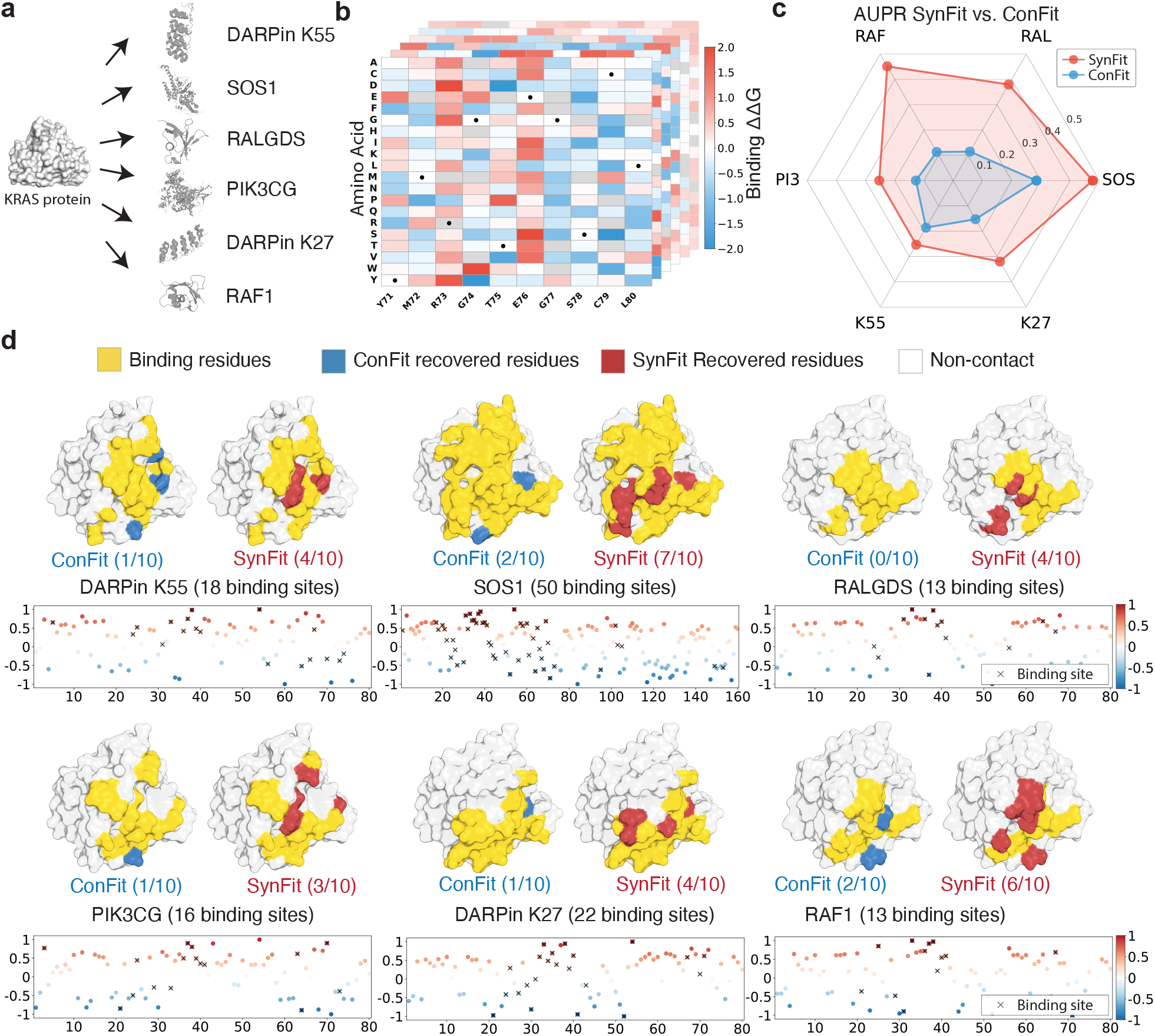
SynFit accurately identifies functional residues across KRAS binding partners by leveraging shared biophysical constraints. **(a)** We curated 6 DMS binding assays from KRAS binding DMS ^33^ corresponding to 6 binding partners. **(b)** After preprocessing, we obtained 6 DMS datasets consisting of 2,418 variants each. (Black dots indicate the wild-type amino acid at each position) **(c)** SynFit achieves higher AUPR performance than individual ConFit model in identification of KRAS binding residues. **(d)** SynFit (bottom panel) performance indicates a substantial improvement compared to individual ConFit models (top panel) in both classification accuracy and positive predictive power for binding sites identification. Yellow highlights indicate partner-specific interface residues; red highlights mark residues contacting all six partners.

We observed that the joint SynFit model consistently outperformed single-assay ConFit models in identifying functional residues, achieving higher AUPR across all binding partners (Figs. 4c,d). Multi-assay training exposed shared functional determinants, such as switch-region and nucleotide-pocket residues, enabling SynFit to disentangle stability-mediated effects from genuine functional disruptions. These results demonstrate that SynFit can uncover shared biophysical constraints governing multi-partner interactions, enabling systematic identification of residues with shared functional roles across multiple binding partners.

Beyond predictive performance, we examined the biological relevance of residues identified as shared across all six binding partners (Fig. 4d, red). These residues partially overlap with known regulatory regions of KRAS, including the switch I (residues 30–38) and switch II (residues 59–67) regions ^35;36^, which mediate conformational changes and effector recognition.

Together, these results show that SynFit not only improves predictive accuracy but also reveals biologically important residues associated with shared functional mechanisms across interaction contexts, providing a valuable method for understanding mutational effects and guiding multi-objective protein design.

### 2.5 SynFit designs combinatorial enzymes with simultaneously optimized activity and enantioselectivity for new-to-nature biocatalysis

We developed a two-stage machine learning workflow that integrates computational library design with multi-objective fitness prediction for biocatalyst optimization (Fig. 5a). First, MODIFY ^37^ designs a starting library balancing evolutionary plausibility with sequence diversity across six active-site positions. After experimental characterization of yield and enantioselectivity, SynFit is trained on these sequence–fitness pairs to predict both objectives across the combinatorial sequence space, and non-dominated sorting identifies Pareto-optimal designs for validation. To experimentally validate SynFit’s predictive capability for use in biocatalysis and enzyme engineering, we applied it to the combinatorial optimization of enzyme variants for biocatalytic asymmetric borylation using N-heterocyclic carbene-boranes (NHC · BH_3_) and *α*-diazocarbonyl compounds, a recently discovered enzymatic reaction ^38^ operating through a carbene transfer mechanism not present in the biological world (Figs. 5b,c). This borylation reaction converts readily accessible building blocks into chiral organoboron products with broad applications in medicinal chemistry and functional materials. We implemented SynFit to optimize biocatalysts derived from the wild-type *Rhodothermus marinus* cytochrome *c* (*Rma* cyt *c*), a compact heme protein consisting of 124 amino acid residues from a hyperthermophile. In optimizing enzyme variants for this reaction, two complementary objectives, overall catalytic yield and enantioselectivity, need to be simultaneously optimized, making it an excellent testing ground for evaluating whether SynFit can jointly model and optimize multiple functional properties. Guided by the crystal structure of *Rma* cyt *c* (PDB ID: 6CUK), we focused our efforts to generate highly active and enantioselective variants by optimizing amino acid residues 75 and 99–103 proximal to heme cofactor in the active site. We first trained SynFit on the 909 experimentally characterized hextuple variants generated by our previously developed method MODIFY ^37^, an unsupervised ML–guided evolution-inspired starting-library design method that balances expected library fitness with sequence diversity. We constructed three non-overlapping train-validation splits and trained multiple SynFit models under different random seeds to ensemble their predictions (Methods). The aggregated predictions were then subjected to non-dominated sorting (Supplementary Note A.1) to identify Pareto-optimal variants (Fig. 5d). The top 100 SynFit-designed variants were produced as an oligo-pool encoding the diversified sequence region, assembled into full-length *Rma* cyt *c* expression constructs in pET-22b(+), and expressed in *Escherichia coli*. The activity and enantioselectivity of these SynFit-designed cytochrome c variants were assessed in whole-cell assays for the enantioselective C–B bond forming reaction.

**Figure 5:**
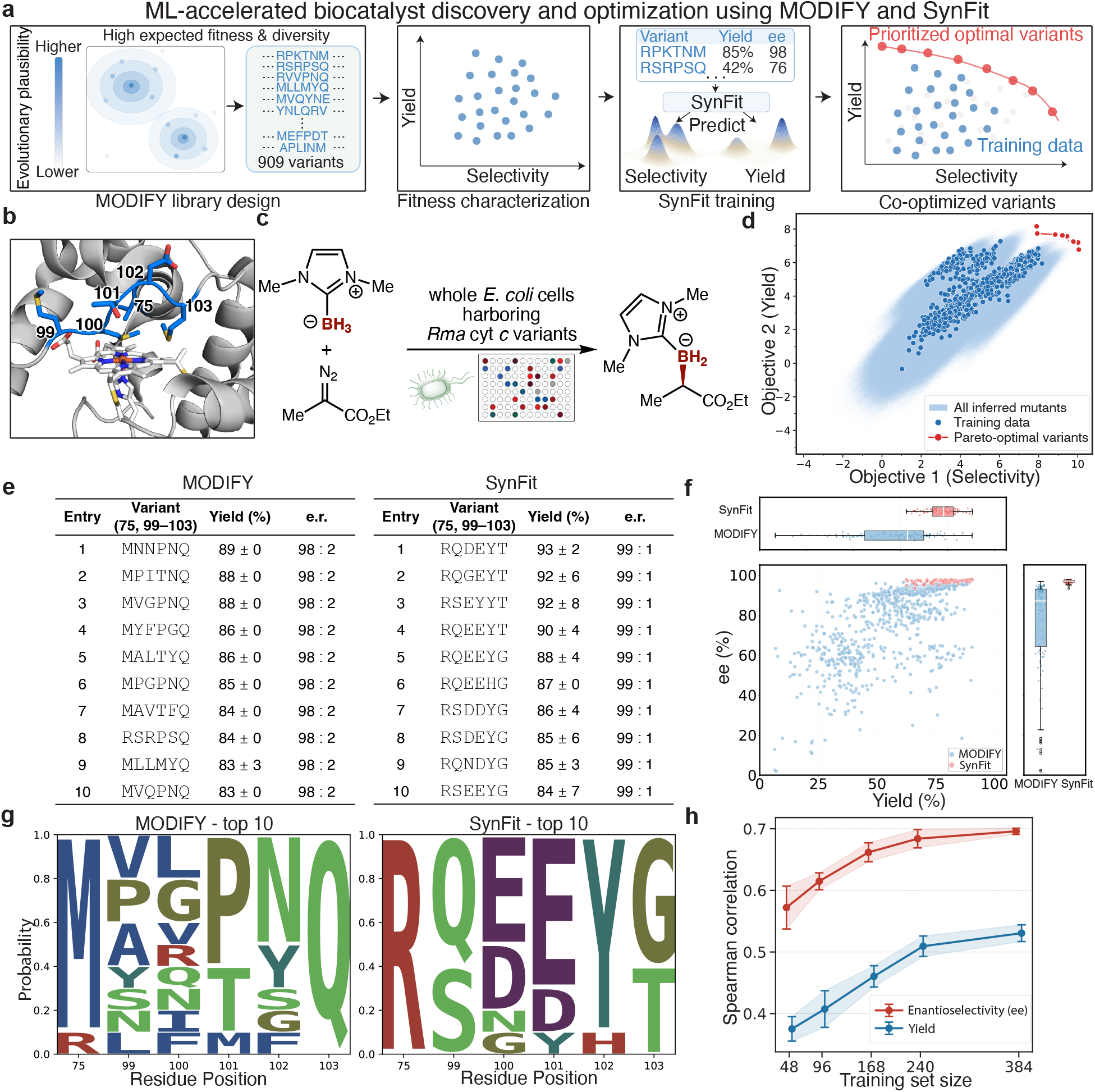
Experimental validation of SynFit for asymmetric carbene B–H insertion catalyzed by *Rma* cyt *c* variants. **(a)** Two-stage biocatalyst design workflow combining MODIFY library design with SynFit multi-objective optimization. **(b)** Structure of *Rhodothermus marinus* cytochrome *c* (PDB 6CUK) highlighting the six targeted active-site residues for combinatorial mutagenesis (positions 75, 99–103). **(c)** Asymmetric carbon–boron bond forming reaction catalyzed by *Rma* cyt *c* variants. Reaction conditions: *E. coli* cells overexpressing *Rma* cyt *c* in M9-N buffer (OD_600_ = 30, ca. 0.03–0.05 mol% *Rma* cyt *c*), NHC-BH_3_ (10 mM), ethyl diazoacetate (10 mM), 5% MeCN, 680 rpm, 12 h. **(d)** SynFit-predicted multi-objective landscape for the 909 experimentally characterized variants. Red points indicate Pareto-optimal *Rma* cyt *c* variants selected for experimental validation. **(e)** Comparison of the ten highest-yielding highly selective variants from the MODIFY and SynFit libraries. Top variants were defined within the highly selective regime and ranked by yield. **(f)** Experimental validation of the top 100 SynFit-predicted variants relative to the original MODIFY-derived dataset, highlighting enrichment of SynFit variants in the high-yield, high-selectivity region. **(g)** Sequence-logo comparison of top ten variants discovered from MODIFY (left) and SynFit (right) libraries exhibiting the highest activity and enantioselectivity, showing stronger sequence convergence among SynFit top variants across the six engineered positions. **(h)** Spearman correlation between predicted and experimentally measured enantioselectivity and yield as a function of training set size for *Rma* cyt *c*.

Starting from the custom-synthesized oligo-pool library encoding SynFit-designed hextuple mutants, we carried out high-throughput biocatalytic borylation reaction using four 96-well plates. Each clone from the 96-well screening was verified by Sanger sequencing. For SynFit variants that appeared more than once in the screening library, catalytic activity and enantioselectivity data were consolidated and standard deviations were calculated to ensure the reliability of our high-throughput results. Together, this effort led to a new biocatalyst activity and enantioselectivity dataset covering a total of 83 variants out of the 100 designs. Across this experimentally validated dataset, SynFit-guided designs exhibited a clear shift toward simultaneously and substantially enhanced yields and enantioselectivities in the multidimensional fitness landscape (Fig. 5f), with a multitude of variants exhibiting superior catalytic performance than all MODIFY variants present in the training data. Unlike MODIFY, which was designed to construct evolution-inspired starting libraries enriched for fitness and sequence diversity, SynFit employs a supervised multi-objective optimization framework that quantitatively links sequence to both catalytic activity and enantioselectivity, enabling the discovery of *Rma* cyt *c* variants with superior catalytic performance beyond the starting library.

Beyond the overall shift in the library-level fitness landscape, we also examined the upper range of practically useful biocatalyst performance among the experimentally validated SynFit variants. Because a range of top-performing *Rma* cyt *c* SynFit variants displayed excellent enantioselectivities up to 99:1 e.r., we ranked these highly enantioselective SynFit variants primarily by yield. It was found that the top 10 SynFit variants all reached yield greater than 84% with an enantioselectivity of 99:1 e.r. (Fig. 5e). Thus, SynFit discovered a range of combinatorial borylation biocatalysts with catalytic performance similar to the experimentally evolved *Rma* cyt *c* resulting from three rounds of site-saturation mutagenesis and screening ^38^.

We next asked whether these top-performing SynFit variants also shared common sequence features. Sequence analysis of the experimentally validated top variants revealed that the principal difference between the two libraries was not only in performance, but also in sequence convergence across the six engineered active-site positions. Notably, SynFit recapitulates key beneficial features of the experimentally evolved variant ^38^, including R75 and T103, while prioritizing a carboxylate side chain (glutamate (E) and aspartate (D)) at residue 100. Wild-type *Rma* cyt *c* features a methionine (M) at residue 100 as a chelator of the heme cofactor. Thus, the preference for E and D at residue 100 reveals a distinctive feature enriched in top-performing borylation biocatalysts, which is also in accord with the experimentally evolved variant ^38^. More interestingly, SynFit also uncovers additional convergent preferences, including Q99, E101, and Y102 at these sites, which diverge from the experimentally evolved variant. In contrast, the top variants from MODIFY were substantially more heterogeneous at these residues (Fig. 5g). This result suggests that SynFit does not merely retrieve isolated high-performing variants, but instead focuses the combinatorial landscape toward a narrower set of residue combinations associated with jointly improved catalytic activity and stereoselectivity. Together, these findings show that SynFit transforms generative, evolution-inspired starting libraries into multi-objective-optimized design spaces, enabling the discovery of new-to-nature biocatalysts with improved activity and selectivity.

In real-world enzyme engineering efforts, due to the limited throughput in chromography-based activity and stereoselectivity screening, experimental datasets are often of the size of 10^2^ variants, making it critical for predictive machine learning models to perform effectively with small training sets. To evaluate the utility of SynFit in biocatalysis research, we examined its performance in data-limited regimes. We performed a low-N scaling analysis by subsampling the training set of *Rma* cyt *c* variants to sizes ranging from 48 to 384 variants (Fig. 5h). SynFit was trained independently on each subset and evaluated on held-out variants using Spearman correlation for both yield and enantioselectivity. Notably, SynFit retained strong predictive accuracy even when trained on fewer than 100 labeled variants, while performance improved steadily as additional data became available. These results highlight the practical utility of SynFit for guiding enzyme engineering in laboratory settings with modestly sized enzyme activity and stereoselectivity datasets.

## 3 Discussion

Protein function is inherently multi-dimensional. A mutation can simultaneously affect stability, activity, binding, expression, or selectivity, and these effects are often mechanistically coupled rather than independent. This coupling is particularly important in protein engineering, where desirable variants rarely optimize only one property in isolation. In many application domains, particularly in biocatalysis and enzyme engineering, the need to simultaneously optimize multiple functional properties, most prominently catalytic activity and selectivity, represents a pervasive and central challenge. These objectives are often interdependent, making the identification of optimal variants difficult using approaches that treat each property independently.

In this work, we present SynFit, a multi-objective protein fitness prediction framework designed to model this interconnected nature of protein function. Our results show that explicitly capturing relationships among functional properties improves the prediction of multidimensional mutation effects and the identification of variants that co-optimize multiple objectives. A central insight of SynFit is that multiple functional assays measured on the same protein contain complementary views of a shared underlying sequence-function landscape. Mutations that disrupt folding can broadly affect abundance, binding, or catalytic activity, whereas mutations at functional sites may selectively impact specific properties. By jointly learning across related objectives, SynFit captures both common constraints critical across multiple functions and objective-specific effects that shape individual functional readouts. The consistent improvements over independently trained models suggest that these cross-assay relationships are biologically meaningful and can be effectively exploited for protein fitness prediction.

This perspective further underscores the importance of multi-objective fitness modeling in guiding protein engineering. In practical settings, the goal is rarely to optimize a single metric, but rather to identify variants that balance multiple functional requirements. This is particularly important in biocatalysis, where achieving high catalytic turnover while furnishing excellent chemo-, regio- and stereoselectivity remains a fundamental challenge. Our benchmark results show that SynFit more effectively prioritizes such variants, indicating that multi-objective learning provides a more holistic representation of protein fitness landscapes than single-property approaches.

The experimental validation in the *Rma* cyt *c* borylation system provides an excellent test case for this capability. In this new-to-nature enzymatic reaction, catalytic activity and enantioselectivity must be simultaneously optimized, reflecting a common constraint in biocatalyst development. SynFit enabled the discovery of a diverse set of cytochrome *c* variants, including numerous sextuple mutants, which exhibit substantial and coordinated improvements in both catalytic activity and stereoselectivity. Notably, these variants populate the high-performance region of the multidimensional fitness landscape and achieve levels of activity and selectivity rivaling or superior than those obtained through multiple rounds of experimental directed evolution. These results highlight SynFit as an effective strategy for navigating complex combinatorial sequence spaces and for identifying practically useful biocatalysts under realistic multi-objective constraints.

Another key aspect of SynFit is its integration of evolutionary priors from pretrained protein language models (pLMs) with assay-specific experimental measurements. Pretrained pLMs capture broad patterns of evolutionary sequence plausibility but do not directly encode functional objectives. SynFit adapts these general representations into function-specific predictive models using experimental fitness data, while preserving their generalizable feature. This enables robust performance even when labeled data are limited, a common scenario in enzyme engineering where throughput constraints restrict dataset size. Together, these design choices make SynFit particularly well suited for real-world multi-objective protein engineering tasks.

Beyond predictive performance, SynFit also offers insights into the molecular mechanisms underlying protein function. When trained across related assays, the model can highlight residues whose mutation effects are consistent across functional measurements, revealing their shared structural or mechanistic constraints. This was illustrated in the KRAS case study, where joint modeling across binding assays enhanced the identification of residues with functional importance across multiple binding partners. Such analyses suggest that multi-objective learning can help distinguish broadly important functional sites from assay-specific effects, extending SynFit’s utility from variant effect prediction to mechanistic interpretation.

Several opportunities remain for further development. Many experimental datasets contain only partial overlap across measured properties, and extending SynFit to more flexibly handle partially overlapped data would broaden its applicability. In addition, while the current SynFit framework relies primarily on sequence-based representations, many functional trade-offs are mediated by structural context and conformational dynamics. Incorporating richer structural information may further improve the modeling of coupled biochemical properties.

Overall, our results support a broader view of protein fitness prediction: multi-assay measurements that characterize different functions of the same protein can be combined to learn more informative fitness prediction models. By enabling the identification of variants that simultaneously improve multiple properties, SynFit provides a general framework for addressing a longstanding challenge in protein engineering. As high-throughput mutational data continue to expand across assays and properties, we envision SynFit as a useful tool for mechanistic studies of protein function and ML-guided protein engineering.

## 4 Methods

### 4.1 Protein Language Models for Protein Fitness Prediction

Recent advances in large-scale protein language models (pLMs) ^39–42^ have demonstrated that amino acid sequences can be modeled analogously to natural language. These models are trained in an unsupervised fashion on massive databases of natural sequences such as UniProt ^43^. Most state-of-the-art pLMs, including ESM3^41^, ProGen3^44^, and ProGen2^13^ adopt a masked language modeling (MLM) objective in which the model is trained to predict masked residues *x*_*i*_ given the surrounding context *x*_*™i*_. Formally, the training objective maximizes the conditional log-likelihood ℒ_MLM_ = Σ_*i*∈𝒮_ log *p*_*θ*_(*x*_*i*_ | *x*_*™i*_), where 𝒮 denotes the set of masked positions. Through pretraining, pLMs capture patterns of residue co-occurrence that reflect evolutionary constraints on protein sequences, providing a notion of how plausible a given mutation is within the space of natural proteins. As a result, zero-shot mutational scoring from pLMs has been shown to correlate with DMS measurements by quantifying the log-likelihood difference between wild-type and mutant residues. However, such unsupervised training does not incorporate function-specific information, which often depends on subtle biochemical determinants beyond evolutionary plausibility. This motivates supervised adaptation strategies, such as ConFit ^23^ and ProteinNPT ^32^, to steer pretrained pLMs toward protein-specific functional objectives. Because these models are trained on large-scale evolutionary data, previous research ^45;46^ has demonstrated the log-likelihood ratio between a wild-type residue and its mutant counterpart provides a useful proxy for relative fitness. Formally, for a mutation 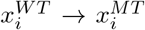, the pLM-derived score is

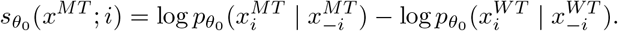

Aggregating across sites yields a zero-shot estimate of mutant plausibility.

Traditionally, supervised fine-tuning approaches such as ConFit ^23^ and ProteinNPT ^32^ focus on optimizing a single functional objective *y* ∈ ℝ (e.g., binding affinity or catalytic activity). Formally, given training data 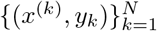, the goal is to learn a mapping *f* : 𝒳^*L*^ → ℝ, *ŷ* (*x*) ≈ *y*, where *x* is a protein sequence and *y* is its corresponding functional measurement. However, in many practical protein engineering settings, multiple properties are simultaneously relevant. For instance, a mutation that improves catalytic yield may reduce thermostability, while a destabilized protein often fails to fold correctly and thus cannot carry out its biological function. Therefore, we are interested in learning predictive models that can jointly optimize multiple objectives and capture their trade-offs. We denote the set of *M* quantitative objectives as 𝒴 = {*y*^(1)^, *y*^(2)^, …, *y*^(*M*)^}, *y*^(*m*)^ ∈ ℝ. Given supervised training data now consists of 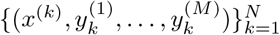, where each *x*^(*k*)^ is a variant of the wild-type sequence and 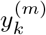 is its experimentally measured fitness under objective *m*. Our aim is to learn a multi-objective model 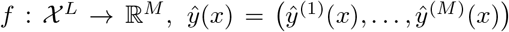, such that each *ŷ*^(*m*)^(*x*) provides an accurate prediction of the true experimental measurement *y*^(*m*)^. By sharing a common representation across tasks while retaining task-specific prediction heads, SynFit leverages correlations between objectives to improve predictive accuracy and robustness.

### 4.2 Overall Training Pipeline

#### Multi-fitness benchmark

We curated a multi-fitness benchmark from ProteinGym ^16^, retaining proteins with multiple functional assays to enable cross-objective evaluation. Datasets with insufficient variant coverage or sequences exceeding GPU memory limits were excluded. The final benchmark comprises 46 DMS assays across 20 proteins, spanning diverse molecular functions including binding, stability, catalysis, and expression.

#### Stage-wise training

To balance task specialization and cross-objective generalization, SynFit employs a novel two-stage training procedure: *Independent head training:* Each task-specific head Head_*m*_ is first trained separately on its corresponding deep mutational scanning (DMS) dataset, while keeping both the pretrained encoder *f*_*θ*_ and the shared module *g*_*ϕ*_ frozen. This ensures that each head captures the fitness signal specific to its objective without interference from other tasks. *Joint fine-tuning:* Once initialized, the heads are frozen and the shared module *g*_*ϕ*_ is trained jointly across all tasks. Gradients from all objectives flow into *g*_*ϕ*_, encouraging it to extract representations that are informative across correlated objectives.

#### Contrastive learning with BT loss and KL regularization

For each objective *m*, we adopt the BT model ^47^, which encourages correct ranking of variants. Given all pairs (*x*_*i*_, *x*_*j*_) such that 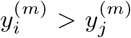, the contrastive loss is

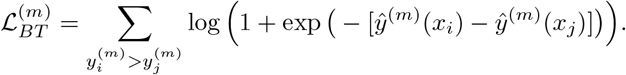

This formulation converts *N* labeled variants into 𝒪(*N*^2^) comparisons, making efficient use of limited DMS data. To prevent the fine-tuned model from diverging too far from the pretrained distribution, we introduce a KL penalty between the predicted amino acid distribution of each head and that of the frozen pLM on the wild-type sequence *x*^*WT*^. Let *p*_pre_(· | *x*^*WT*^) be the residue distribution from the pretrained model and 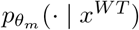 the distribution induced by head *m*. The KL regularizer is

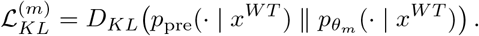

This regularization stabilizes training by anchoring predictions to the evolutionary prior encoded in the pretrained model, reducing overfitting to small-scale experimental datasets.

#### Per-task and overall objective

The training objective for each task combines contrastive supervision and KL regularization:

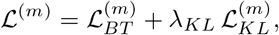

where *λ*_*KL*_ controls the strength of regularization. The global multi-task loss aggregates across objectives:

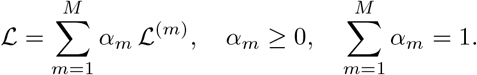

In practice, *α*_*m*_ can be set to equal weights (*α*_*m*_ = 1/*M*) or tuned to emphasize specific objectives.

### 4.3 Function Site Identification

We adapt the methodology of our previous SPURS model ^26^ to identify functionally important sites, replacing ESM1v sequence-likelihood scores with the task-specific prediction heads of SynFit, while retaining SPURS-predicted ΔΔ*G* as the stability reference signal. For a wild-type protein sequence **x**^WT^ and a single-residue mutation 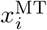 at position *i*, SynFit produces task-specific prediction scores

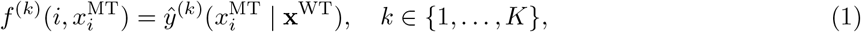

where *K* = 6 for the six KRAS binding partners and 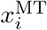 is the substituted amino acid. These scores serve as our proxy for the relative functional fitness of each single-mutation variant under binding partner *k*. We use SPURS ^26^ to predict the folding free-energy change ΔΔ*G* for every single-residue substitution and define a per-variant stability score as the clip-and-normalised negative SPURS-predicted ΔΔ*G*,

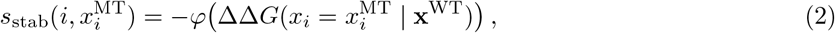

where *φ*(·) clips values to the 0.1^th^ and 99.9^th^ percentiles of the ΔΔ*G* distribution and rescales to [0, 1], so that stabilizing mutations receive high scores and destabilizing ones receive low scores. To disentangle stability effects from binding-specific contributions, we fit a sigmoid function mapping SynFit prediction scores to SPURS stability scores across all single-mutation variants of each protein:

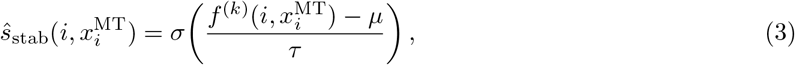

where *µ* and *τ* are location and scale parameters estimated by weighted least squares and *σ*(·) denotes the sigmoid function. The deviation of the observed stability score from the sigmoid prediction,

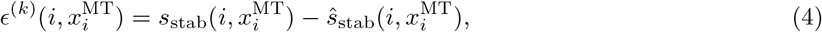

indicates whether a mutation’s stability effect is larger or smaller than expected from its functional score ^48^. We summarise per-site importance as the *function score*: the mean residual across all substitutions at residue *i*,

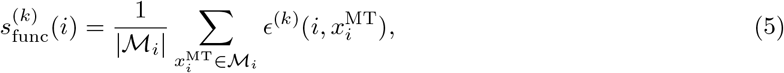

where ℳ_*i*_ is the set of all 19 non-wildtype amino acid substitutions at position *i*.

To curate our training dataset, we removed variants containing stop codons or higher order mutations, and retained only variants with complete measurements across all 6 binding assays, resulting in a total of 2,418 variants. During training, we employed a three–fold data-splitting strategy to ensure robust model evaluation and complete coverage of the dataset. For each fold, 90% of the data were randomly assigned to the training set and the remaining 10% were held out as the validation set. Three independent models were trained on these non-overlapping splits, such that each data point appeared in the validation set exactly once across the three folds. This design ensures that the combined set of validation predictions covers the entire dataset without redundancy.

While individually trained ConFit models achieve strong performance on their respective DMS assays, SynFit further improves upon these single-task models. By jointly training SynFit on all 6 binding DMS datasets across all binding partners (DARPin K55 (PDB ID: 5O2T, SOS1 (PDB ID: 1NVW), RALGDS (PDB ID: 1LFD), PIK3CG (PDB ID: 1HE8), DARPin K27 (PDB ID: 5O2S), and RAF1 (PDB ID: 6VJJ).), SynFit leverages mutual information shared across assays - for example, the critical common contact residues that contribute to multiple binding interfaces. This joint training enables SynFit to refine the function scores 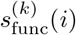 and more accurately identify residues consistently under multi-partner functional constraint. The fact that SynFit outperforms all six ConFit baselines demonstrates that it effectively integrates information across shared binding sets, capturing functional signals that are not accessible to individually optimized models.

### 4.4 Application of SynFit to engineering *Rma* cyt *c* biocatalysts for carbene C–B bond formation

SynFit is experimentally validated on biocatalytic C–B bond-formation reaction catalyzed by *Rhodothermus marinus* cytochrome *c* (*Rma* cyt *c*). Building on the previously reported MODIFY-designed ^37^ *Rma* cyt *c* library, we expanded experimental coverage from the initially tested 160 variants to a total of 909 sequence–function pairs. The designed sequences were synthesized as an oligo-pool library, cloned into a pET-22b(+) vector, and co-expressed with the heme-maturation plasmid pEC86 in *E. coli* BL21(DE3). Single colonies were picked into 96-well plates for small-scale expression under identical IPTG and 5-aminolevulinic acid induction at 20 °C. Following expression, cells were collected by centrifugation, resuspended in M9–N buffer, and directly used as whole-cell catalysts for the carbene C–B bond-formation reaction in 96-well plates under anaerobic conditions using N-heterocyclic carbene borane and ethyl diazoacetate as substrates. After 12 h, reactions were quenched and analyzed by chiral HPLC to determine total yield and enantioselectivity (e.r.). In parallel, the variants were characterized by PacBio long-read sequencing to verify sequence identity and ensure accurate sequence–function correspondence across the dataset. This standardized workflow produced a consistent, comparable dataset comprising 468 sequences from the intended MODIFY library and 441 additional recombined or non-designed variants across the same six active-site positions, providing a comprehensive experimental foundation for SynFit training and evaluation.

During training, we employed a three-split validation strategy to ensure both robustness and full dataset coverage. The 909 MODIFY variants were partitioned into three non-overlapping validation subsets, each comprising 10% of the data, while the remaining 90% served as the training set for that split. This ensured that every variant appeared in the validation set exactly once across the three runs. To further enhance robustness and account for stochastic variation, the full three-split procedure was repeated under five independent random seeds, resulting in a total of fifteen trained SynFit models. During inference, we averaged the predicted scores from all models to obtain the ensemble output for both enantioselectivity and yield. These ensemble predictions were ranked using non-dominated (Pareto) sorting, with the mean of both fitness scores resolving ties. (Supp. MethodsA.1) Because exhaustively sorting all possible mutants is not computationally realistic, we first identified a high-confidence candidate set by selecting variants that exceeded the 99.5th percentile of predicted scores in both objectives. All variants meeting this dual-percentile criterion were retained for non-dominated sorting. In this way, the 100 highest-ranking variants were selected for oligo-pool synthesis and experimental characterization.

## Acknowledgments

This research is supported by the National Science Foundation (No. 2442063 and No. 2435754 to Y.L., and ITE-2448848 to Y.Y. on ML methods for directed evolution), the NSF Molecule Maker Lab Institute funded by the NSF under Awards No. 2019897 and 2505932, and the National Institutes of Health (R35GM150890 to Y.L. and R35GM147387 to Y.Y. on metalloenzyme research). Y.Y. is an Alfred P. Sloan Research Fellow (FG-2024-22244), a Camille Dreyfus Teacher-Scholar Awardee (TC-25-084), a David & Lucile Packard Fellow (2023-76169) and a Howard Hughes Medical Institute Freeman Hrabowski Scholar. Computations were carried out at the Advanced Cyberinfra-structure Coordination Ecosystem: Services & Support (ACCESS) Program, supported by NSF awards (CHE-260005 to Y.Y.). This work used computational resources provided by the CloudHub GenAI Seed Grant funded by GaTech IDEaS and Microsoft. We thank members of the Luo Lab at Georgia Institute of Technology and the Yang Lab at the University of California Santa Barbara for helpful discussions.

## Data, Materials, and Software Availability

All code and data necessary to reproduce the results in this study are publicly available at: https://github.com/luo-group/SynFit. Additional materials are available from the corresponding author upon reasonable request.

## Supplementary Materials

### Supplementary Methods

#### A.1 Non-dominated Sorting

The complete selection pipeline integrates four complementary algorithms to identify optimal protein mutations from a large candidate space. First, **Algorithm 1** (Non-Dominated Sorting) orchestrates the hierarchical ranking process by repeatedly calling **Algorithm 2** (ExtractNonDominatedSet) to partition solutions into Pareto fronts, where each front represents a distinct level of multi-objective optimality. The domination relationships are determined by **Algorithm 3** (Dominates), which implements the formal definition of Pareto dominance under maximization objectives—ensuring that solutions in earlier fronts are superior trade-offs between competing objectives. Finally, **Algorithm 4** (SelectTopK) addresses the practical constraint of selecting a fixed number of variants for experimental validation by including complete high-quality fronts and applying an average-score tie-breaking criterion on the boundary front. This ensures the final selection maintains both Pareto optimality (by prioritizing better fronts) and internal quality (by selecting the best candidates within the partial front), producing a diverse yet high-performing set of mutations that span the Pareto frontier while respecting experimental budget constraints. In practice, we utilized the PyGMO library’s fast non-dominated sorting implementation, which employs efficient data structures to achieve *O*(*MN* ^2^) complexity for the complete sorting process.

##### Algorithm 1 Non-Dominated Sorting for Multi-Objective Optimization. This algorithm partitions a population of solutions into hierarchical Pareto fronts based on dominance relationships. It iteratively extracts non-dominated solutions to form each front, where Front 1 contains globally optimal solutions (the Pareto frontier), Front 2 contains solutions dominated only by Front 1, until all solutions are assigned a rank

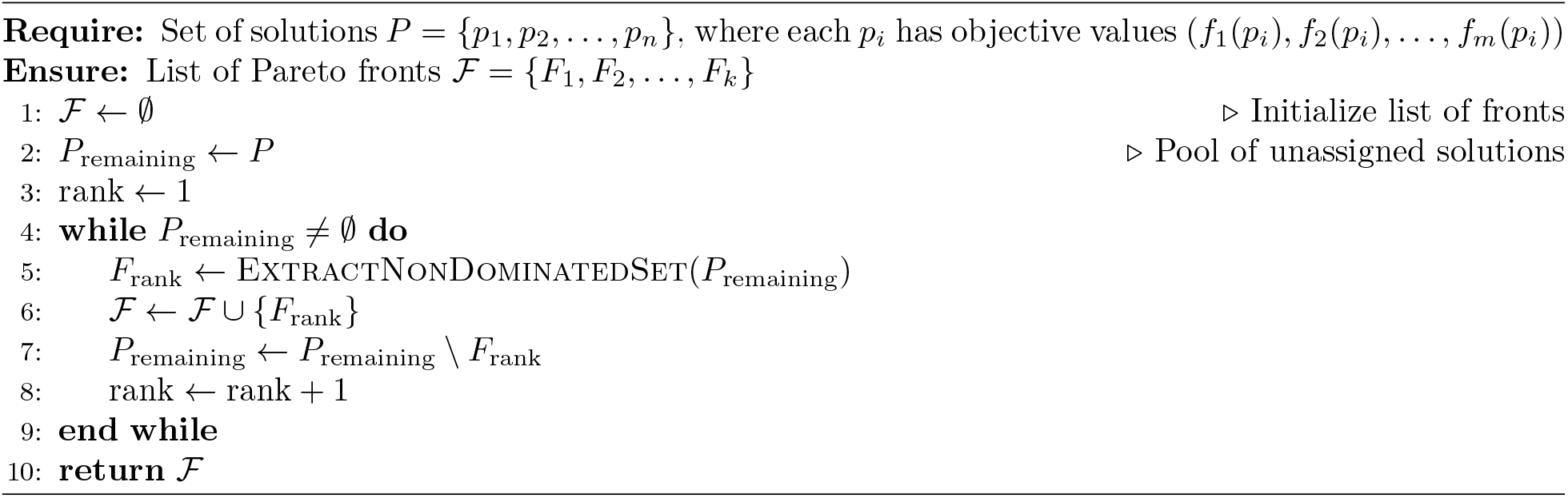

##### Algorithm 2 ExtractNonDominatedSet. This subroutine identifies the set of non-dominated solutions. For each candidate solution, it checks whether any other solution dominates it; if no dominating solution exists, the candidate is added to the non-dominated set. This forms the basis for constructing each Pareto front in the sorting process

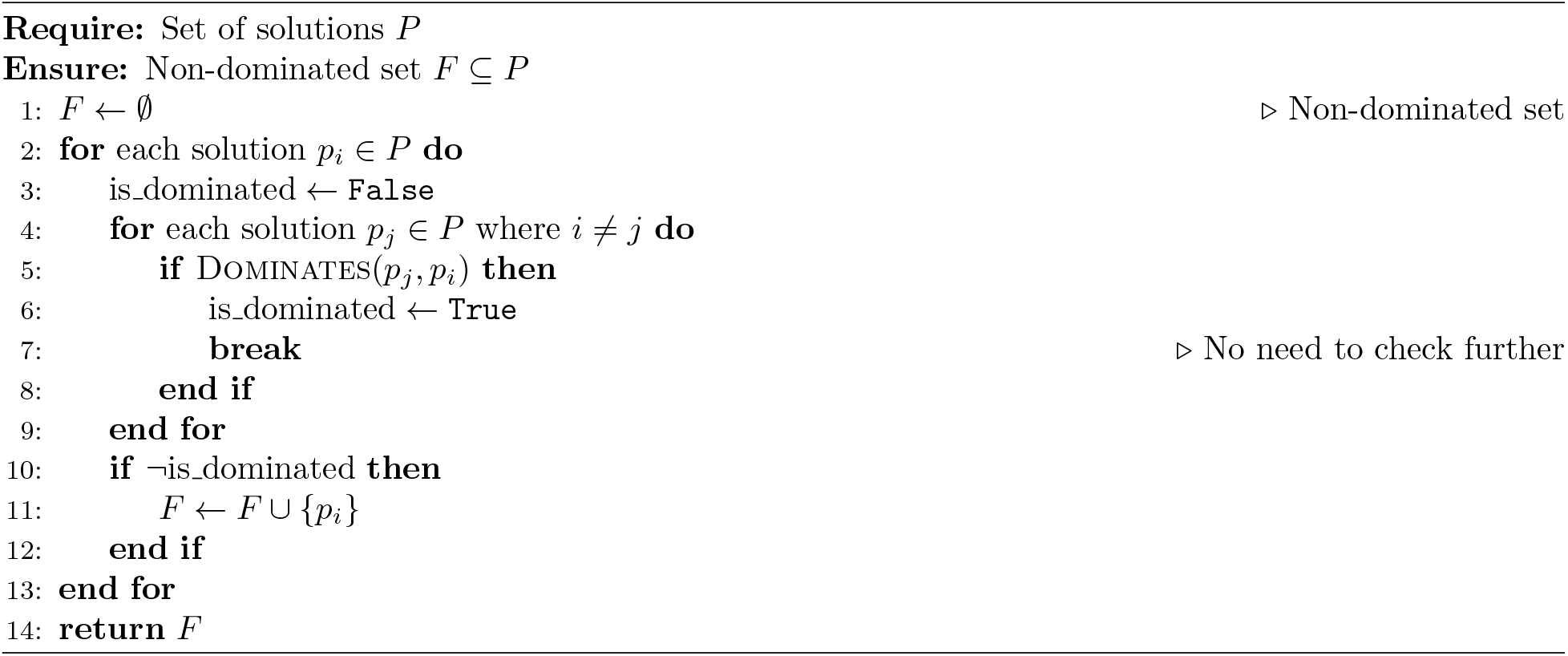

##### Algorithm 3 Dominates (Maximization). This function determines whether solution *p*_*i*_ dominates solution *p*_*j*_ under a maximization objective framework. Dominance occurs when *p*_*i*_ is at least as good as *p*_*j*_ in all objectives and strictly better in at least one objective. The function returns true only when both conditions are satisfied, enabling proper Pareto frontier identification

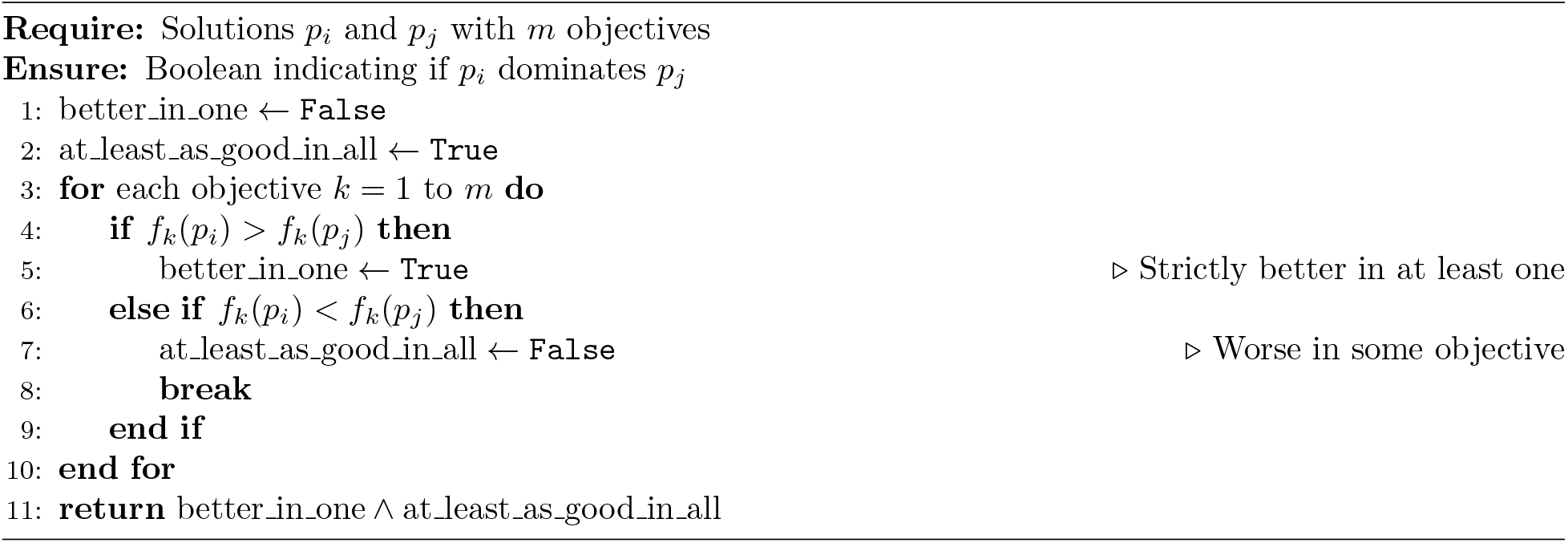

##### Algorithm 4 SelectTopK with Tie-Breaking. This algorithm selects exactly *K* solutions from the ranked Pareto fronts for experimental validation. It greedily includes complete fronts sequentially until adding the next front would exceed *K*. For the final partial front, it breaks ties by ranking solutions based on their average objective score and selecting the top-ranked solutions to reach exactly *K* total, ensuring both quality and diversity in the final selection

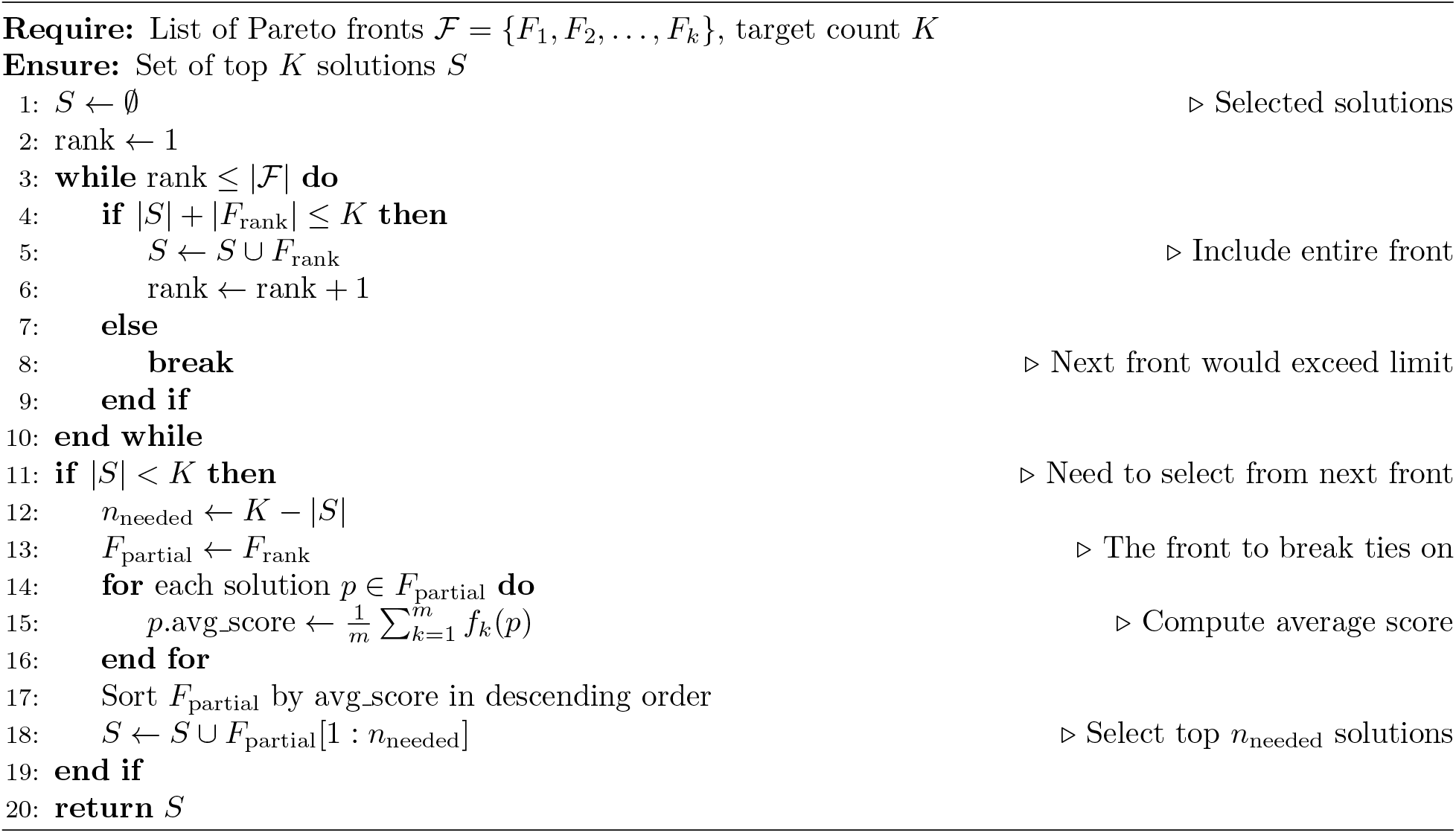

### B Supplementary Tables

#### B.1 Multi-fitness benchmark datasets

We initially collected 22 proteins encompassing 50 deep mutational scanning (DMS) datasets from ProteinGym ^16^, focusing on proteins for which multiple functional assays were available. To ensure sufficient coverage and computational feasibility, we excluded the NPC1 HUMAN dataset ^49^, which contained only 58 variants and was too small for reliable benchmarking, and omitted SPIKE SARS2^50^ due to its long sequence length exceeding GPU memory limits during model training. After filtering, our final benchmark comprised 20 proteins with 46 functional assays spanning diverse reaction types and molecular functions, forming a large-scale multi-fitness dataset for evaluating cross-objective generalization in SynFit. In section 2.3, due to the fact that some DMS assays measure overlapping functions or lack wild-type references, our analysis was limited to eight proteins – *CP2C9 HUMAN, HXK4 HUMAN, KCNE1 HUMAN, RL40A YEAST, KCNJ2 MOUSE, OXDA RHOTO, PTEN HUMAN*, and *RASK HUMAN* – spanning 16 assays in total.

**Table S1:** Summary of the multi-fitness benchmark curated from ProteinGym, containing 46 DMS assays across 20 proteins with diverse functional measurements such as binding, expression, and growth rates, supporting comprehensive evaluation of SynFit models.

| UniProt ID | #Variants | Fitness Type(s) |
| --- | --- | --- |
| SPIKE_SARS2 | 3,798 | Binding<sup>50</sup>, Expression<sup>50</sup> |
| KCNE1_HUMAN | 2,313 | Expression<sup>51</sup>, Activity<sup>51</sup> |
| Q53Z42_HUMAN | 3,345 | Binding<sup>52</sup>, Expression<sup>52</sup> |
| VKOR1_HUMAN | 693 | Expression<sup>53</sup>, Activity<sup>53</sup> |
| A0A2Z5U3Z0_9INFA | 2,351 | Growth Rate<sup>54</sup>, Growth Rate<sup>55</sup> |
| RL40A_YEAST | 1,139 | Growth Rate<sup>56</sup>, Growth Rate<sup>57</sup>, Activity<sup>58</sup> |
| OXDA_RHOTO | 6,388 | Activity<sup>59</sup>, Expression<sup>59</sup> |
| CAR11_HUMAN | 2,355 | Growth Rate<sup>60</sup>, Growth Rate<sup>60</sup> |
| CCDB_ECOLI | 1,150 | Activity<sup>61</sup>, Growth Rate<sup>61</sup> |
| HXK4_HUMAN | 8,256 | Growth Rate<sup>62</sup>, Expression<sup>63</sup> |
| KCNJ2_MOUSE | 6,790 | Activity<sup>64</sup>, Expression<sup>64</sup> |
| S22A1_HUMAN | 9,716 | Expression<sup>65</sup>, Activity<sup>65</sup> |
| BLAT_ECOLX | 989 | Growth Rate<sup>66</sup>, Growth Rate<sup>67</sup>, Growth Rate<sup>68</sup>,<br>Growth Rate<sup>69</sup> |
| RPC1_LAMBD | 352 | Activity<sup>70</sup>, Activity<sup>70</sup> |
| DYR_ECOLI | 2,362 | Growth Rate<sup>71</sup>, Growth Rate<sup>72</sup> |
| P53_HUMAN | 1,049 | Growth Rate<sup>73</sup>, Growth Rate<sup>73</sup>, Growth Rate<sup>73</sup>,<br>Growth Rate<sup>74</sup> |
| RASK_HUMAN | 23,073 | Expression<sup>33</sup>, Binding<sup>33</sup> |
| SPG1_STRSG | 1,459 | Binding<sup>75</sup>, Binding<sup>76</sup> |
| CP2C9_HUMAN | 4,422 | Expression<sup>77</sup>, Activity<sup>77</sup> |
| PTEN_HUMAN | 4,840 | Expression<sup>78</sup>, Activity<sup>79</sup> |
| SRC_HUMAN | 3,357 | Activity<sup>80</sup>, Activity<sup>81</sup>, Growth Rate<sup>82</sup> |

### C Supplementary Figures

#### C.1 Complete multi-objective pareto analysis results

**Figure S1:**
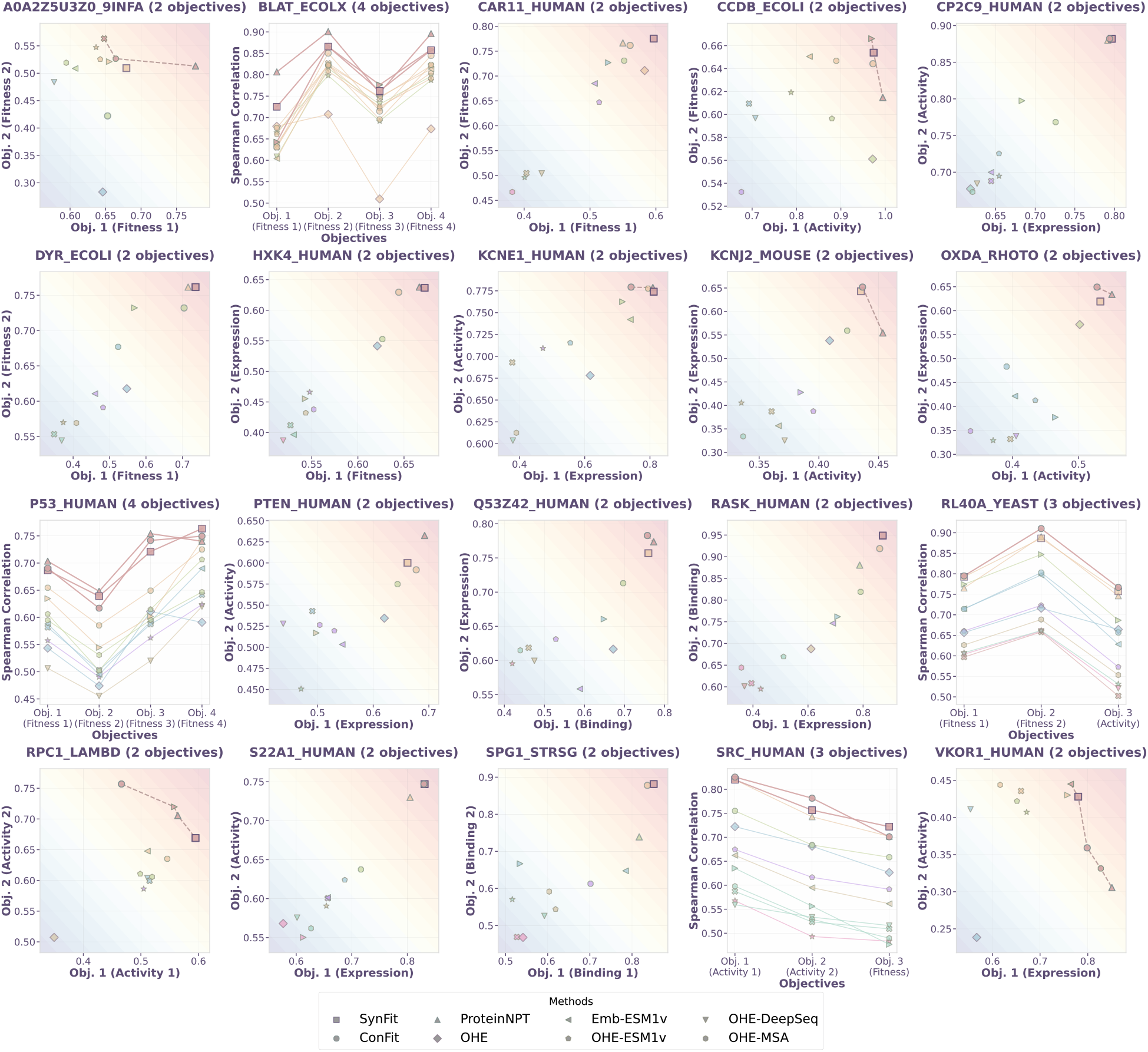
Multi-objective protein Pareto analysis. Comprehensive Pareto analysis across the first subset of proteins in the multi-fitness dataset. Each panel summarizes trade-offs between distinct fitness objectives (e.g., activity, stability, expression, binding) for SynFit and baseline models, with Pareto fronts highlighting the advantage of multi-objective joint training.

#### C.2 Functional Site Identification

**Figure S2:**
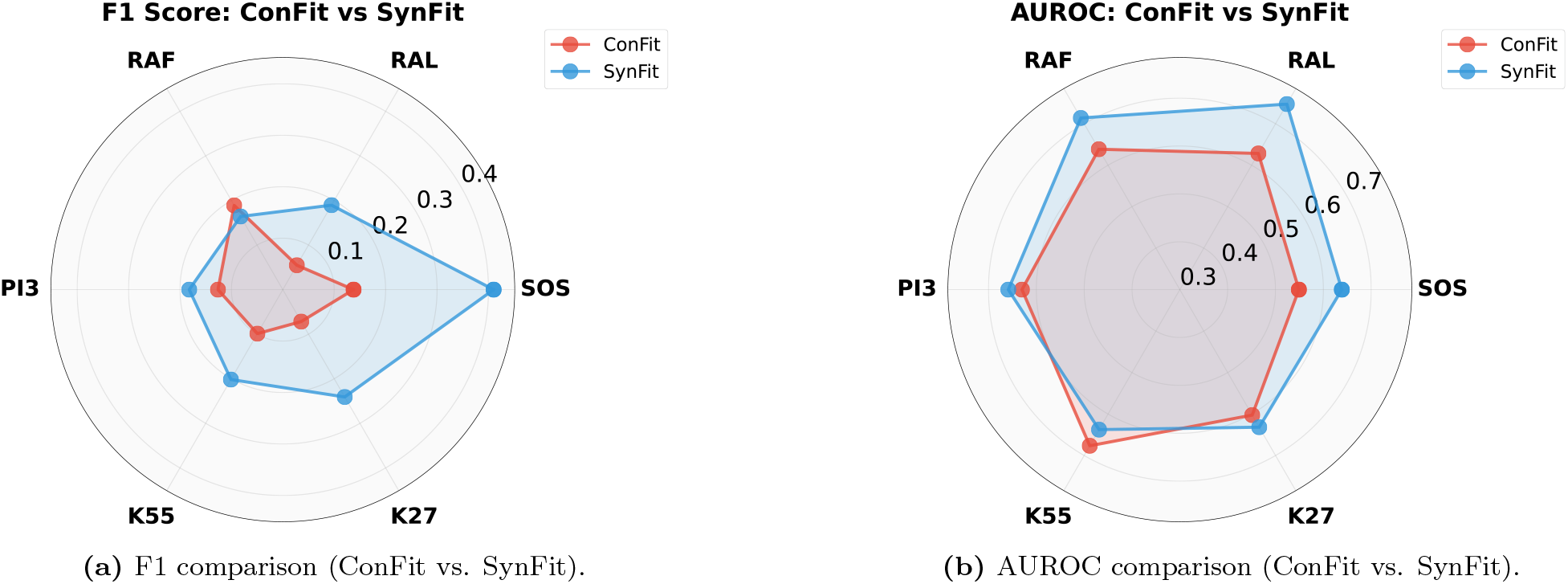
Function site identification SynFit vs. ConFit AUROC and F1 score performance. Side-by-side comparison of F1 and AUROC across six KRAS binding partners. SynFit consistently outperforms ConFit on both metrics on five out of the binders for functional site identification, indicating improved classification performance.

## Notes

### Competing Interest Statement

The authors have declared no competing interest.

